# Iodine staining as a useful probe for distinguishing amyloid fibril polymorphs

**DOI:** 10.1101/2020.03.21.001214

**Authors:** Takato Hiramatsu, Naoki Yamamoto, Seongmin Ha, Yuki Masuda, Mitsuru Yasuda, Mika Ishigaki, Keisuke Yuzu, Yukihiro Ozaki, Eri Chatani

## Abstract

It is recently suggested that amyloid polymorphism, i.e., structural diversity of amyloid fibrils, has a deep relationship with pathology. However, its prompt recognition is almost halted due to insufficiency of analytical methods for detecting polymorphism of amyloid fibrils sensitively and quickly. Here, we propose that iodine staining, a historically known reaction that was firstly found by Virchow, can be used as a method for distinguishing amyloid polymorphs. When insulin fibrils were prepared and iodine-stained, they exhibited different colors depending on polymorphs. Each of the colors was inherited to daughter fibrils by seeding reactions. The colors were fundamentally represented as a sum of three absorption bands in visible region between 400-750 nm, and the bands showed different titration curves against iodine, suggesting that there are three specific iodine binding sites. The analysis of resonance Raman spectra and polarization microscope suggested that several polyiodide ions composed of I_3_^−^ and/or I_5_^−^ were formed on the grooves or the edges of β-sheets. It was concluded that the polyiodide species and conformations formed are sensitive to surface structure of amyloid fibrils, and the resultant differences in color will be useful for detecting polymorphism in a wide range of diagnostic samples.

## Introduction

Amyloid fibrils are protein aggregates associated with many amyloidosis and neurodegenerative diseases including Alzheimer’s disease (AD) and Parkinson’s disease^1,2^. While straight unbranched morphology and cross-β structure are known as a common basic structure of amyloid fibrils, it was revealed that amyloid fibrils show structural variations even from the same protein; this phenomenon is referred to as amyloid polymorphism^3–5^. Variations in the number and the arrangement of protofilaments are an example of polymorphism^6,7^, and more microscopic variations at the level of secondary or tertiary structure of polypeptide chains have been also identified in many proteins^8–14^. In studies of prion diseases, different physicochemical properties among amyloid polymorphs, such as fragility and growth rates, have been suggested to be the molecular basis underlying the presence and propagation of distinct prion strains^15^. Amyloid polymorphs also exert different neurotoxic effects on living cells^9^, and it has recently been suggested that tau protein adopted distinct amyloid conformations in the human brain in different diseases^14^. The formation of such conformationally distinct amyloid fibrils now attracts much attention for understanding amyloid-related pathologies and developing curative treatments in terms of protein structure.

Amyloid polymorphs have been successfully reproduced in vitro depending on growth conditions, e.g., temperature ^15^, protein concentration^16^, and solvent^17^; however, the most critical factors have not been fully revealed yet^5^. Subtle variations in the amino acid sequence among different species also result in different fibril structures^18^, although it does not correspond to amyloid polymorphism because amino acid sequences are not the same. In another case, distinct fibril structures can be also induced in a stochastic manner even under the same conditions^19^. Recently, detailed structural investigations of amyloid fibrils are progressing by using several powerful methods, such as solid-state NMR (ssNMR)^9,11^, cryo-electron microscopy (cryo-EM)^12–14^, and X-ray crystallography^8^. The clarified structures have revealed distinct molecular structures of amyloid polymorphs. The relevance of amyloid polymorphism to in vivo pathology has also been clarified by the structural investigation of Aβ_1-40_ amyloid fibrils that were prepared with seeds derived from brain tissues of AD patients with distinct clinical histories^20^. However, the use of these techniques requires a lot of labor and time and therefore making them unsuitable for high throughput use. If an easy-to-use method of analyzing fibril structures can be developed, it will become a great help to cover a wide range of samples including pathological tissues as targets of investigations, thereby contributing to progress toward a unified understanding of relationships between fibril structures and pathologies.

Here, we focus on iodine as a molecular probe that recognizes structural polymorphs of amyloid fibrils. Iodine staining is a coloring reaction that occurs by complexing iodine with crystalline compounds or polymers. The molecular origin of the color is the formation and subsequent alignment and orientation of polyiodide ions. An iodine-starch complex is a good example, and from the crystal structure of iodine-doped α-cyclodextrin, it is suggested that linear chains of pentaiodide ions (I_5_^−^) are encapsulated in channel-like voids in the center of the helical polysaccharides^21,22^. The coloration by polyiodide ions is also observed in iodine-doped synthetic polymers such as polyacetylene^23^, nylon 6^24^ and polyvinyl alcohol^25^. Interestingly, colors formed by iodine staining vary sensitively depending on constituting polyiodide ions and their coordination patterns inside the complex structures^26,27^. From these observations, it is postulated that the color exhibited by iodine staining will serve as a sensitive indicator of structures of host compounds to which iodine binds. With regard to iodine staining of amyloid fibrils, Virchow firstly found that tissue deposits of amyloid fibrils were stained with iodine^28^. However, soon after this pioneering discovery in the 1850s, fluorescent dyes such as Congo Red and thioflavin S or thioflavin T (ThT) with higher detection sensitivity replaced iodine for diagnosis of amyloidosis^29–31^, and molecular mechanisms underlying iodine reaction have almost not been studied; indeed, there is only one report by Dzwolak^10^.

In this work, we have performed iodine staining of human insulin amyloid fibrils using three polymorphs obtained under different additive conditions (i.e., in the presence of NaCl or SDS and in the absence of additives). Insulin is a hormone protein associated with iatrogenic amyloidosis^2^. It has been also used as a good model for amyloid researches in a past few decades^32^. As a result of the characterization of color properties of each type of insulin fibrils by ultraviolet-visible (UV-Vis) spectra, we have found that the color significantly varied depending on the polymorphs. Each color conserved even after the self-seeding reaction, demonstrating that iodine staining tracked the propagation of amyloid polymorphs. Polyiodide ions formed by iodine staining were characterized by resonance Raman spectra and titration experiments. Furthermore, iodine-stained insulin spherulites were observed by polarization microscopy to gain insights into the binding manner of the polyiodide ions to amyloid fibrils. It was proposed that different polyiodide species were formed depending on the surface structures of amyloid fibrils, which resulted in different colors. From the results obtained, molecular mechanisms of iodine staining of amyloid fibrils and future perspectives for the use of iodine staining as a probe for amyloid polymorphism will be discussed.

## Results

### Formation of polymorphic insulin amyloid fibrils

We firstly prepared amyloid fibrils of human insulin used for iodine staining. In this work, 25 mM HCl and 65 °C were selected as standard conditions by referring to previous reports^33–35^, and amyloid polymorphs were attempted to be obtained by using two types of additives, NaCl and SDS. As a result of incubating the sample solutions for 24 hours, ThT fluorescence exhibited different intensity among the amyloid fibrils formed (Fig. 1a, blue bars). In light of a previous observation that the affinity and stoichiometry between ThT and amyloid fibrils varied depending on fibril structures^36^, the different fluorescence intensity suggests that amyloid polymorphs were formed. Indeed, when seed-dependent fibril formation was performed with the amyloid fibrils formed to produce daughter fibrils, relative differences in ThT fluorescence intensity was conserved (Fig. 1a, red bars). The daughter fibrils all exhibited higher intensity than that of the parent fibrils, probably because non-fibrillar aggregates that could not self-propagate contained in the parent fibrils. Furthermore, significant differences in peak wavelength were found in addition to the differences in intensity in ThT fluorescence spectra (Fig. 1b), which strongly supports the formation of amyloid polymorphs. We refer to the amyloid fibrils generated in the absence and presence of NaCl and SDS as no-salt fibrils, NaCl fibrils, and SDS fibrils, respectively.

**Figure 1.**
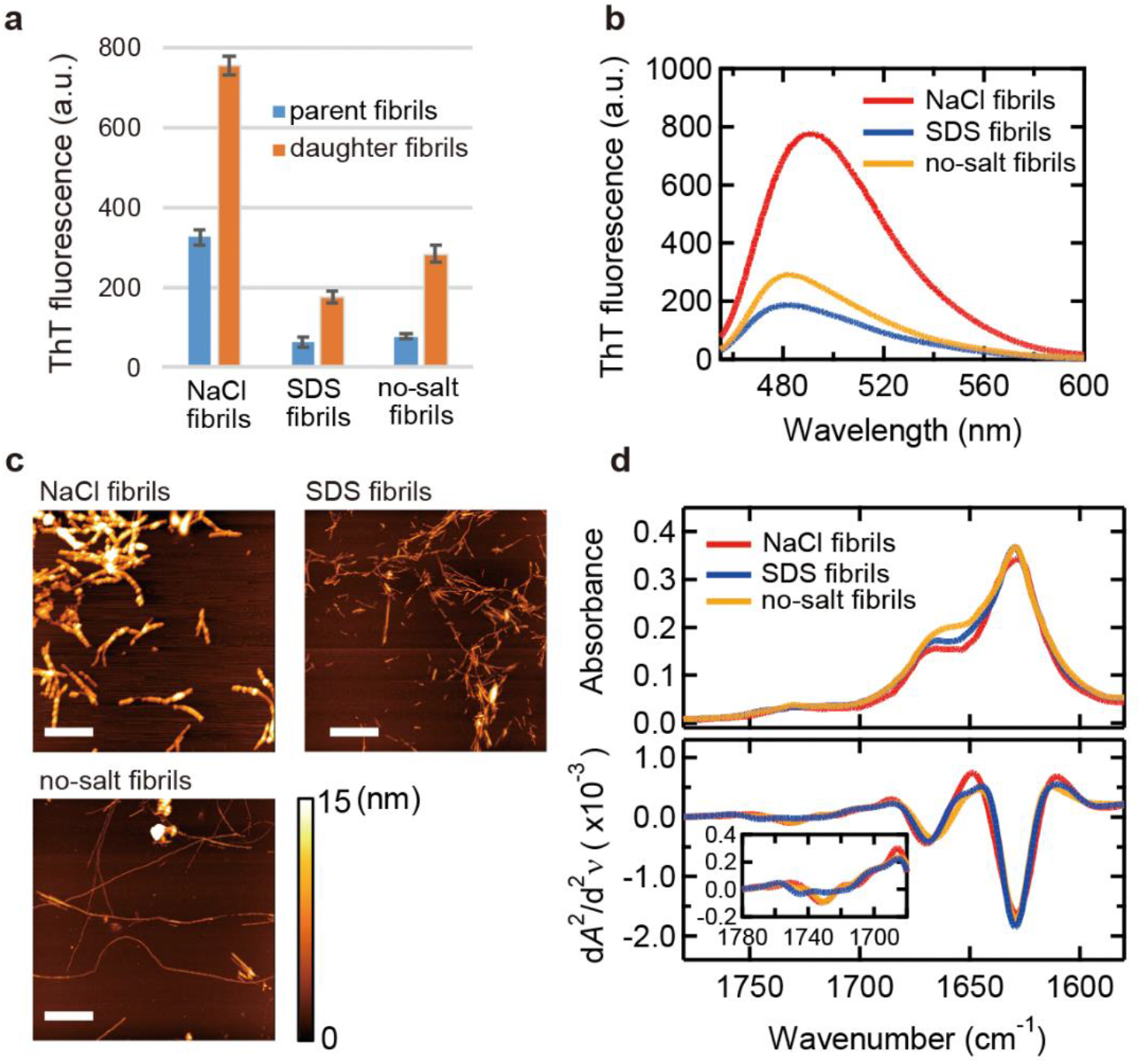
Basic properties of three types of insulin amyloid fibrils prepared in the presence of NaCl (NaCl fibrils) or SDS (SDS fibrils) or in its absence (no-salt fibrils). (**a**) ThT fluorescence intensity. Blue and orange bars represent fluorescence intensity of parent and daughter fibrils, respectively. All measurements were performed three times and the error bars depict the S.D. ± mean. (**b**) ThT fluorescence spectra of daughter fibrils. (**c**) AFM images. The white scale bars represent 1 μm. The color scale bar represents the height of samples. (**d**) ATR-FTIR absorption spectra (upper panel) and their second derivatives (lower panel) in amide I region.

We next analyzed morphology and secondary structure of the amyloid fibrils generated from the three different additive conditions by AFM and ATR-FTIR spectroscopy. AFM images showed variety in fibril thickness, and the NaCl fibrils were markedly thicker (~10 nm) compared to SDS (~ 4 nm) and no-salt fibrils (~ 4 nm) (Fig. 1c). On the other hand, when ATR-FTIR spectra were analyzed, all fibrils exhibited a main peak at 1630 cm^−1^ in amide I region, which was assigned to β-sheet structure (Fig. 1d; upper panel)^37^. Second derivatives of the spectra also showed similar peak positions of minima among NaCl, SDS, and no-salt fibrils. However, differences at around 1670 cm^−1^ and 1710-1750 cm^−1^, which are assigned to β-turn and carboxyl groups of side chains, respectively, were observed (Fig. 1d; lower panel)^37^. This result suggests that polymorphism was derived from differences in turn and/or higher-order structures including hierarchical architecture, while the main cross-β structure was very similar.

### Color formation by iodine staining of insulin amyloid fibrils

To investigate iodine staining of amyloid fibrils, we analyzed changes in colors by mixing iodine solution with NaCl, SDS, or no-salt fibrils. The iodine solution, in which three types of iodine species, I^−^, I_2_, and I_3_^−^, are present in equilibrium, shows yellowish color, and the absorption spectrum of the iodine solution shows two major peaks and one broad one derived from I_3_^−^ and I_2_, respectively^38,39^ (Fig. S1). When amyloid fibrils were added to the iodine solution, reddish or bluish color appeared for all types of fibrils (Fig. 2a). Native insulin mixed with iodine solution, on the other hand, showed quite a similar UV-Vis absorption spectrum to that observed for the iodine solution-only control (Fig. S1a), suggesting that iodine staining occurs specifically to fibril structures. Interestingly, the color varied among types of fibrils, and absorption spectra in UV-Vis region showed different shapes in visible region (400-750 nm) (Fig. 2b). The spectra as well as their second derivatives showed that iodine-stained NaCl fibrils had a major peak at around 550 nm (Fig. 2b, red line), whereas iodine-stained SDS and no-salt fibrils had one at around 650 nm (Fig. 2b, blue and orange lines, respectively). Although contamination of salts from the parent fibrils may affect the spectral shape, this was ruled out by confirming that the spectra did not change even after the addition of salts (Fig. S2).

**Figure 2.**
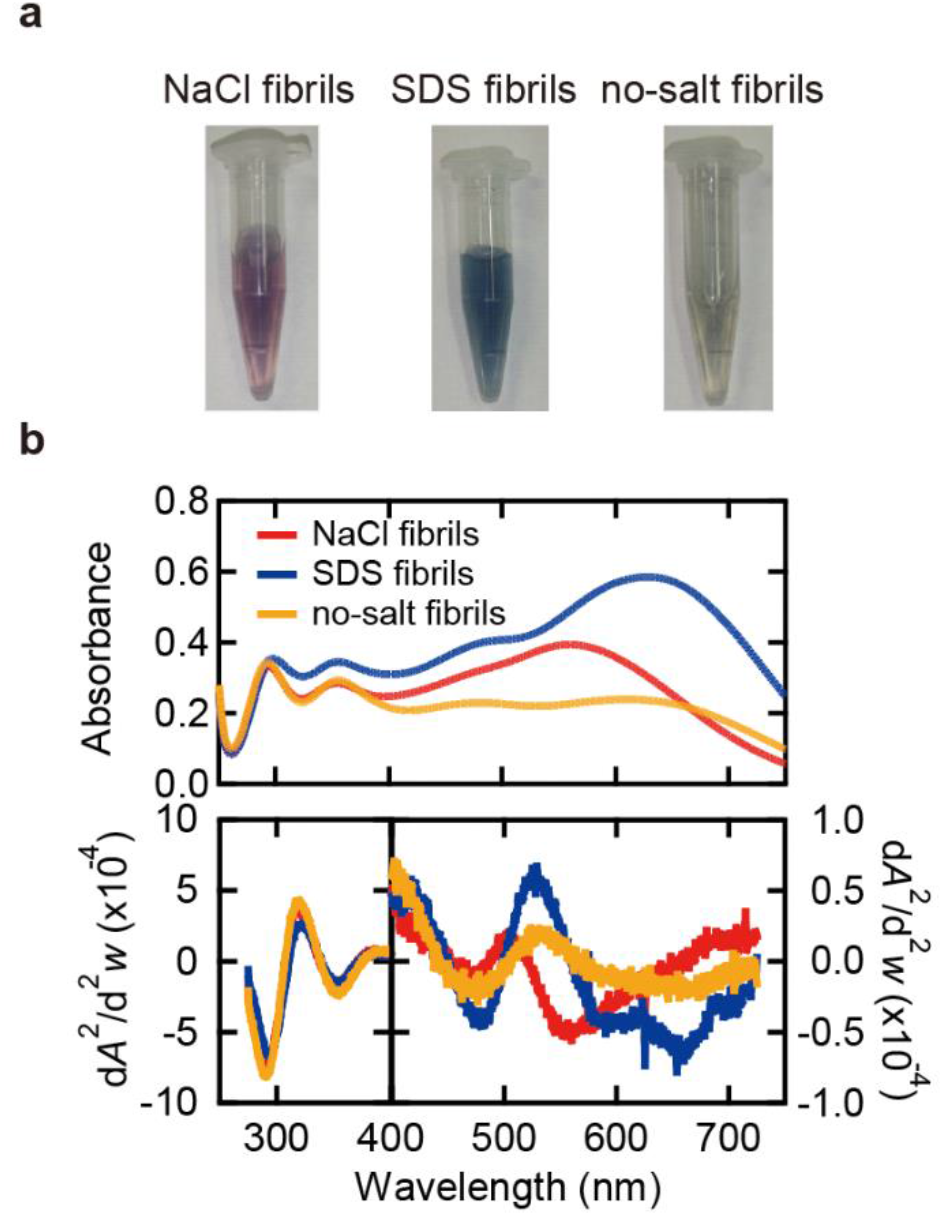
Color properties of iodine-stained insulin amyloid fibrils. (**a**) Photographs of samples of iodine-stained NaCl fibrils, SDS fibrils, and no-salt fibrils. (**b**) Absorption spectra in UV-Vis region (upper panel) and their second derivatives (lower panel) of iodine-stained fibrils. For each absorption spectrum, the spectrum of unstained fibrils was subtracted to obtain net spectrum derived from iodine molecules. It should be noted that the scales of the vertical axes for the region of 400-740 nm and that of 260-400 nm are different in the lower panel. The conditions of iodine staining were 0.25 mg/ml insulin, 0.3 mM KI and 0.04 mM I_2_ in 25 mM HCl.

We next performed spectral deconvolution to discuss absorption bands constructing the experimental spectrum. Each spectrum was replotted against wavenumber so that the analysis was performed in a manner proportional to energy, and then was reproduced by a sum of Gaussian distributions according to the following model function;

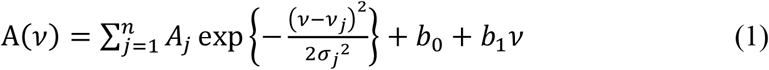

where *A*_*j*_ and *v*_*j*_ represent amplitude, center wavenumber of the *j*th absorption band, respectively, and *σ*_*j*_ is a coefficient related to the full width at half-maximum (FWHM) of the *j*th absorption band, and FWHM is described as 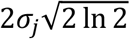. *b*_0_ and *b*_1_ are intercept and slope of the baseline for Rayleigh scattering caused by fibrils, respectively. Curve fitting was performed until both measured and second derivative spectra were reproduced well. For stable and accurate convergence, the wavenumber and FWHM of the most prominent band in visible region were fixed to values read from the local minimum and its distance to the inflection point in the second derivative spectrum.

As a result, all of the absorption spectra were reproduced successfully by approximating with six Gaussians (i.e., *n* = 6) and a linear baseline (Fig. 3). Although hardly recognizable in the second derivative spectrum (see Fig. 2b), the installation of another band between 23 × 10^3^ and 25 × 10^3^ cm^−1^ (i.e., between 400 and 435 nm), which was conceived to be derived from I_2_ molecules (see Fig. S1)^38,39^, certainly improved the reproduction of the experimental spectra. When the six Gaussian absorption bands were compared to the bands observed in the iodine solution (Fig. S1b), the three bands at higher wavenumbers (dashed lines in Fig. 3a-c) showed similar positions to those for the I_2_ and I_3_^−^ in the iodine solution (dashed lines in Fig. S1b). The remaining three absorption bands at lower wavenumbers, on the other hand, were observed only for the fibril-bound iodine (filled bands in Fig. 3a-c). It was thus suggested that the color formation by the iodine staining of amyloid fibrils was represented by these three newly emerged absorption bands, although, in the cases of NaCl and SDS fibrils, the absorption band at ~25 × 10^3^ cm^−1^ was increased after iodine staining and thus is partly responsible for the color formation. The three newly emerged bands showed higher peak wavenumbers in NaCl fibrils compared to those of SDS and no-salt fibrils, and in SDS and no-salt fibrils, the relative intensities of the three bands were different while the peak positions were similar.

**Figure 3.**
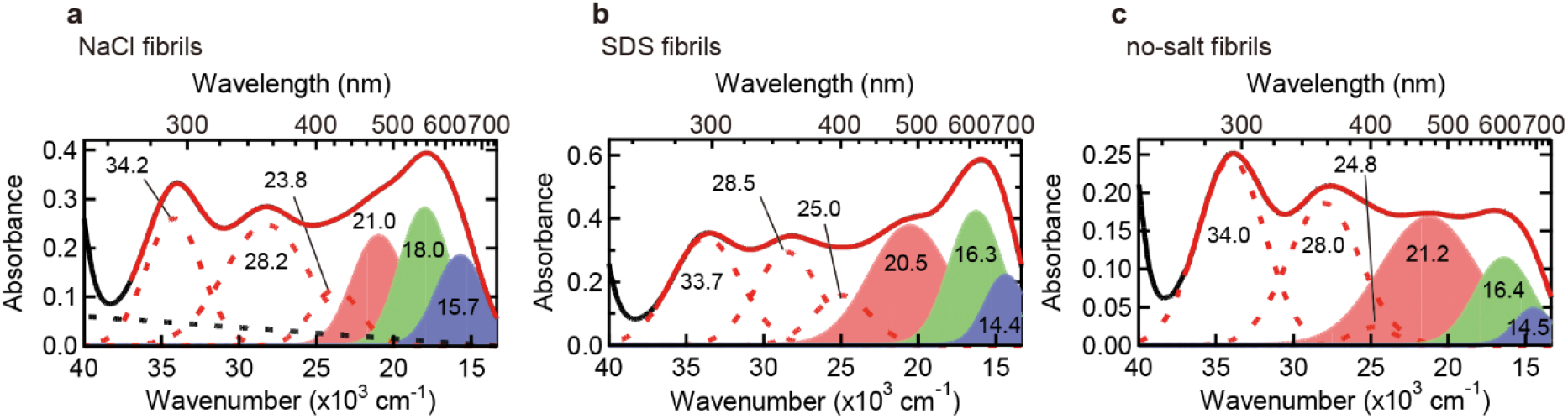
Deconvolution of UV-Vis absorption spectra of iodine-stained NaCl fibrils (a), SDS fibrils (b), and no-salt fibrils (c). In all panels, black and red solid lines represent the experimental and reproduced spectra, respectively, and the experimental spectra are the same as those shown in Fig. 2b. Six Gaussians represented with dashed red lines or filled areas correspond to absorption bands obtained by the deconvolution using eq. 1, and dashed black lines represent baselines derived from scattering of the sample fibrils. The wavenumber of each absorption band is labeled.

### Tracking structural propagation of amyloid fibrils by iodine staining

To verify whether the intrinsic iodine colors can serve as a probe for the propagation of amyloid fibril structures, we tracked time course of seed-dependent elongation by iodine staining (Fig. 4). In this experiment and thereafter, NaCl and SDS fibrils, which showed markedly distinct characteristics in terms of the positions of the three absorption bands, were subjected to the analyses. As seeds, the parent fibrils were used. When the color formation by iodine was examined by sampling the reaction mixture at different time points, absorption intensity in visible region gradually increased both in the NaCl and SDS fibrils as time advanced (Fig. 4a,c). Spectral deconvolution was successfully achieved by assuming six absorption bands with constant wavenumbers (Fig. S3). When the areas of the three absorption bands at lower wavenumbers were plotted against reaction time, all of them increased synchronously with the time-dependent increase of ThT fluorescence intensity (Fig. 4b,d), suggesting that the color developed in response to fibril elongation without changing its tone.

**Figure 4.**
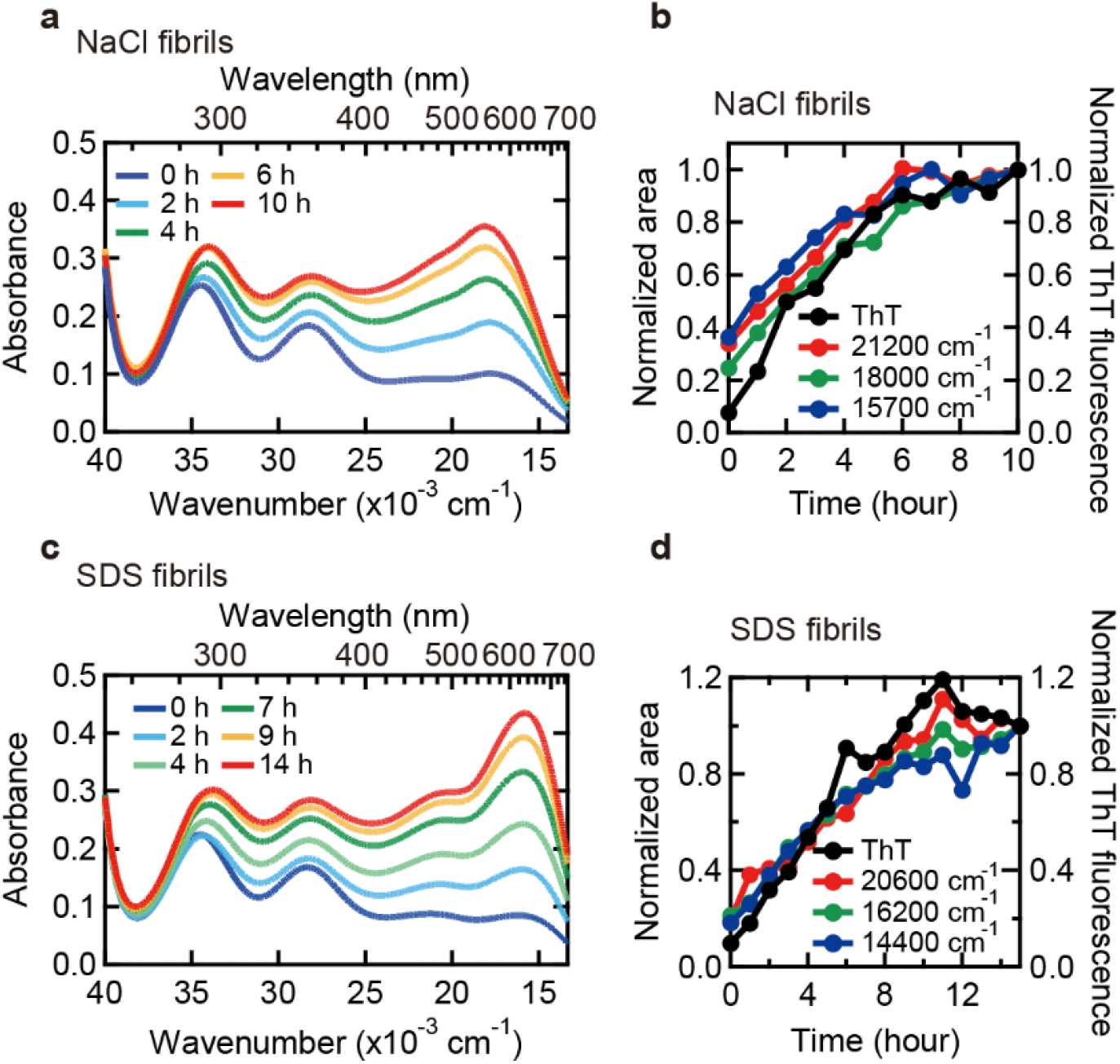
Tracking seed-dependent fibril growth by iodine staining. (**a**-**d**) Time dependent change in iodine color of NaCl (**a**,**b**) and SDS fibrils (**c**,**d**). In panels **a**and **c**, UV-Vis spectra of NaCl and SDS fibrils (**c**) stained with iodine monitored at different time are shown. The conditions of iodine staining were 0.25 mg/ml insulin, 0.3 mM KI and 0.04 mM I_2_ in 25 mM HCl. In panels **b**and **d**, areas of the three absorption bands of NaCl fibrils (**b**) and SDS fibrils (**d**) in the low wavenumber region (e.g., filled bands in Fig. S3c-h) are plotted against reaction time. For details about spectral deconvolution for the determination of these absorption bands, see Fig. S3. ThT fluorescence intensity which was measured concurrently is also plotted for comparison.

### Characterization of polyiodide species responsible for visible color

According to studies of iodine-doped compounds, the molecular mechanism of visible color formation by iodine is due to the formation of polyiodide ions^40,41^. In the case of iodine-doped amyloid fibrils, the formation of pentaiodide ions and several other oligoiodide ions was suggested by Dzwolak^10^. To identify polyiodide species formed on the amyloid fibrils analyzed in the present work, resonance Raman spectra were measured. Raman spectroscopy is a useful technique for detecting subunits of polyiodide chains in complexes with amylose^22,42,43^ and other organic compounds^44–47^. Here, two excitation wavelengths 514.5 nm (19436 cm^−1^) and 785 nm (12739 cm^−1^) were used; two out of the three absorption bands (red and green bands in Fig. 3a-c) are supposed to be excited with the 514.5 nm laser, and the remaining band (blue bands in Fig. 3a-c) is supposed to be excited with the 785 nm laser.

As a result, resonance Raman effects were observed for both NaCl and SDS fibrils, and characteristic Raman spectra with two strong peaks at around 110 cm^−1^ and 160 cm^−1^ were observed at both excitation wavelengths (Fig. 5). Based on a previous Raman study of the crystalline compounds containing triiodide or pentaiodide ions^44^, the 110 cm^−1^ and 160 cm^−1^ peaks were assigned to I_3_^−^ and I_5_^−^, respectively. The present observation therefore suggests that the fundamental building blocks of polyiodide ions are I_3_^−^ and I_5_^−^. Unstained fibrils showed no significant peaks in the same way as those of iodine solution without fibrils (Fig. 5, dotted line), indicating that the peaks are derived from iodine species bound to the fibrils. When two spectra excited at different wavelengths were compared in NaCl or SDS fibrils (i.e., Fig. 5a,c or 5b,d), the positions and intensity ratio of the I_3_^−^ and I_5_^−^ peaks were different. This observation suggests that the conformations and the composition ratio of the I_3_^−^ and I_5_^−^ subunits constituting the polyiodide ions vary among the absorption bands. Especially when excited at 514.5 nm, two weak bands were additionally observed at ~ 220 cm^−1^ and ~ 320 cm^−1^, which were assumed to be overtone bands of the I_3_^−^ and I_5_^−^ bands, respectively. A very weak combination band of the I_3_^−^ and I_5_^−^ bands was also found at around 270 cm^−1^ for SDS fibrils, whereas it was undetectable for NaCl fibrils, probably due to low intensity of the 160 cm^−1^ band. When excited at 785 nm, an overtone band of the I_5_^−^ band as well as a combination band were observed both for NaCl and SDS fibrils. When the spectra of NaCl and SDS fibrils excited at the same wavelength were compared (i.e., Fig. 5a,b or 5c,d), the intensity ratio of the I_3_^−^ and I_5_^−^ peaks was significantly different between NaCl and SDS fibrils at both excitation wavelengths, and the positions of the peaks were also slightly different. This observation implies that different polyiodide ions are formed depending on amyloid polymorphs.

**Figure 5.**
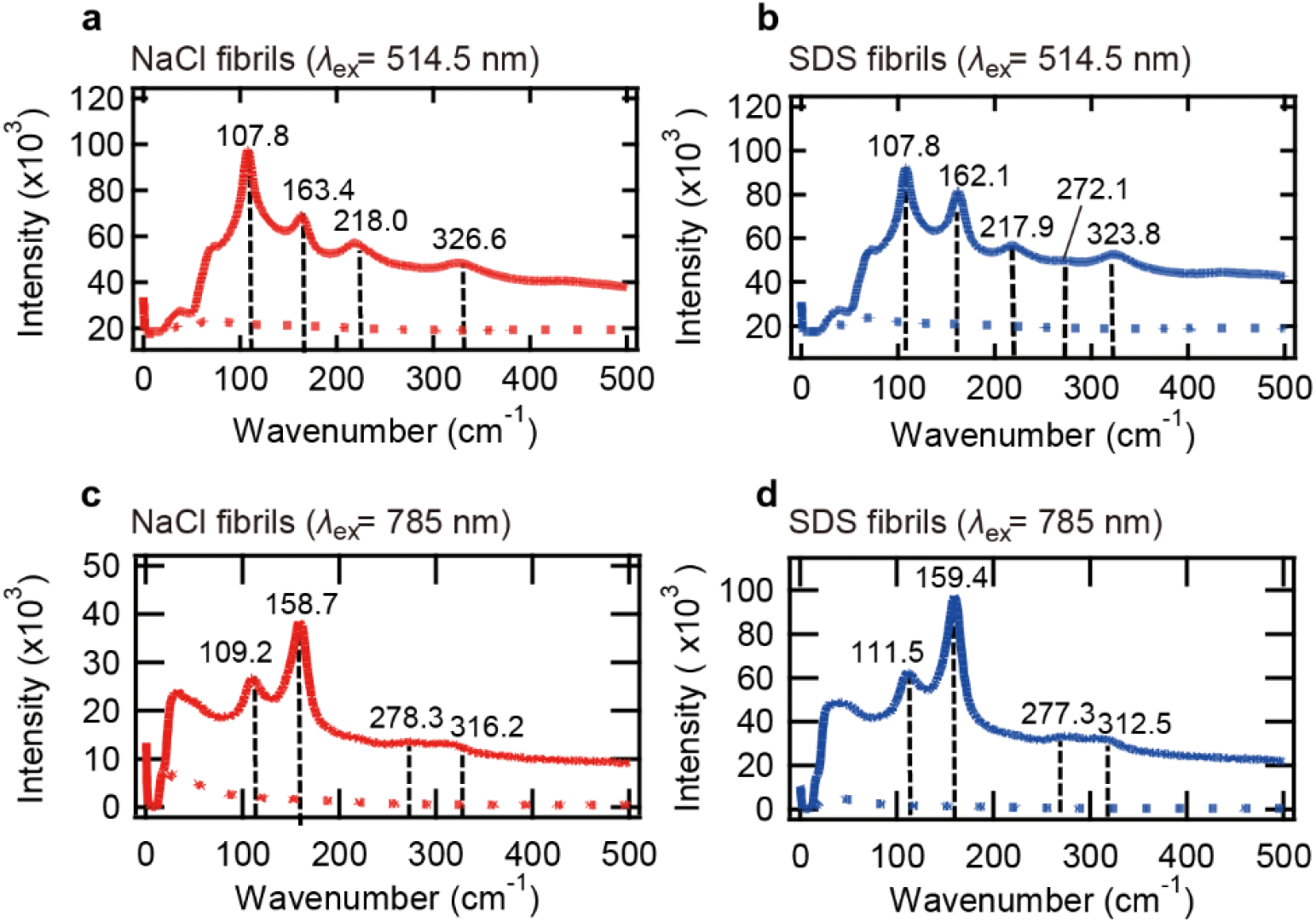
Resonance Raman spectra of iodine-stained insulin amyloid fibrils with excitation of 514.5 nm (a,b) and 785 nm (c,d). The spectra of NaCl fibrils (**a**,**c**) and SDS fibrils (**b**,**d**) are shown. Samples containing 0.5 mg/ml amyloid fibrils, 0.6 mM KI, and 0.08 mM I_2_ in 25 mM HCl were subjected to the measurements. Solid and dotted lines represent spectra of iodine-stained fibrils and unstained fibrils measured as a control, respectively, and the positions of peaks are indicated by black dashed lines.

### Titration of iodine to insulin amyloid fibrils

We further performed titration experiments to iodine concentration dependence of the absorption band intensities. With stepwise addition of the iodine solution to amyloid fibrils, gradual intensification of the absorption bands was observed both in NaCl and SDS fibrils. The absorption in the low wavenumber region finally saturated after the addition of excess amount of iodine (Fig. 6a,b). Interestingly, absorption intensity at lower wavenumbers grew slightly later compared to that at higher wavenumbers, and indeed, distinct shapes of the titration plots were observed when the areas (*A*(*c*)) of the three absorption bands in the low wavenumber region (the bands with filled area in Fig. S4) were plotted against iodine concentration *c* (Fig. 6c,d). The titration plots showed a cooperative properties and as a result of curve fittings with Hill equation^48^,

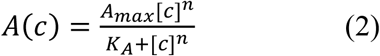

where *A*_max_, *K*_A_, and *n* represent band area attained after saturation, iodine concentration at which half of the binding sites are occupied, and Hill coefficient, respectively, different parameters in their binding affinity and cooperativity were obtained among the three absorption bands both in NaCl (Fig. 6c) and SDS fibrils (Fig. 6d). This result suggests that there are at least three different iodine-binding sites. Furthermore, all plots exhibited *n* > 1, from which a cooperative mechanism was suggested in the iodine binding to amyloid fibrils. It is deduced that small iodide species, such as I_3_^−^ and I^−^ with negligible absorption in visible region initially bind and then facilitate subsequent iodine binding to form polyiodide ions showing visible color.

**Figure 6.**
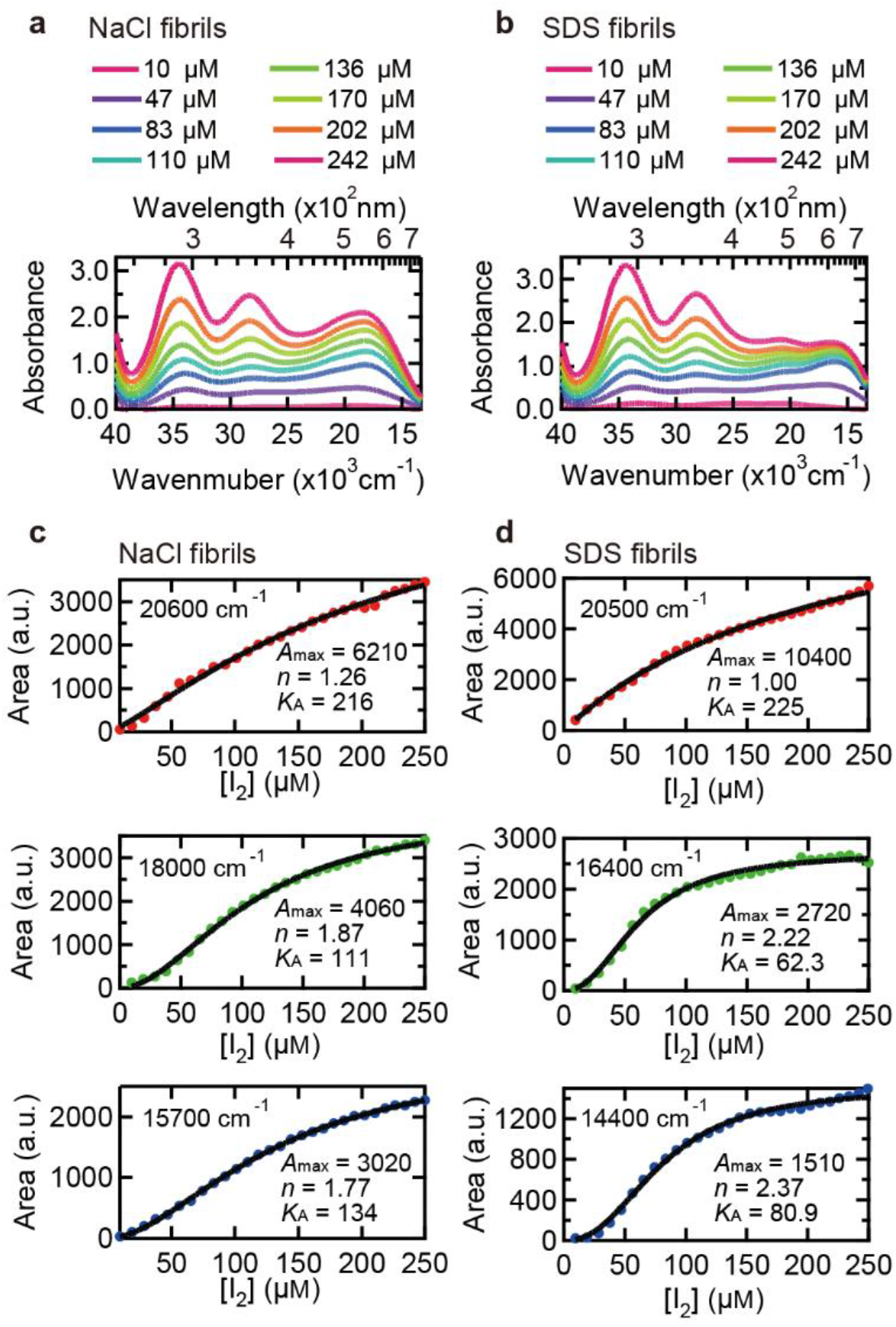
Titration of insulin fibrils with iodine solution. 0.5 mg/ml insulin fibrils in 25 mM HCl and iodine solution containing 18 mM KI and 2.4 mM I_2_ in 25 mM HCl were used as an analyte and a titrant, respectively. (**a**,**b**) Examples of absorption spectra of NaCl (**a**) and SDS fibrils measured at different concentrations of iodine solution. (**c**,**d**) Iodine concentration dependence of area of the three absorption bands of iodine-stained NaCl or SDS appearing in the low wavenumber region (e.g., filled bands in Fig. S4c-h). In these plots, the concentration of I_2_ in the titrant iodine solution is represented as the scale of horizontal axes. For details about spectral deconvolution for the determination of these absorption bands, see Fig. S4. Black lines represent fitted curves using eq. 2 and parameters obtained from the fitting are represented inside the panels.

### Investigation of the orientation of polyiodides on the surface of amyloid fibrils

It is often revealed that polyiodide chains formed by complexation of iodine/iodide with various compounds lie in an oriented manner^21,47,49^. To gain insights into whether the polyiodides bound to amyloid fibrils have some specific orientation, a polarization property of the iodine-stained fibrils was investigated. Linear polyiodides ions have transition moment in a direction parallel to their long axes^44^, and thus the aligned polyiodides ions absorb polarized light, as seen in unidirectionally stretched films of iodine-doped polyvinyl alcohol^50^. In this analysis, amyloid spherulites, a spherical hierarchical structure composed of radially oriented amyloid fibrils^51^, were selected as a good model for evaluating the orientation of fibril-bound iodine molecules. Amyloid spherulites of insulin were formed, treated with iodine solution, and then observed by a microscope equipped with an analyzer to assess whether the transmitted light of the iodine-stained amyloid spherulites showes a polarized property.

As a result, the transmission microscope images of the iodine-stained amyloid spherulites showed two cone-shaped dark sections along the analyzer axis, and their direction changed with response to the rotation of the analyzer (Fig. 7). In contrast, no dark section was observed for unstained spherulites (Fig. S5), verifying that the polarized nature of the iodine-stained spherulites was derived from polyiodide. The direction of the dark sections and that of the analyzer axis coincided well, which suggests that polyiodide chains are oriented along the fibril axis given the radial orientation of fibril axes in the amyloid spherulite.

**Figure 7.**
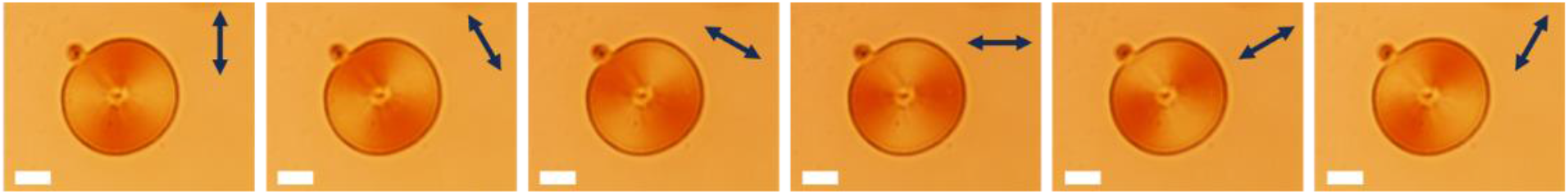
Anisotropic property of polyiodides on the spherulite. In each image, the direction of the analyzer slit is represented with a double-headed arrow in the upper right. Scale bars represent 10 μm. The images of an unstained spherulite recorded by the same procedure are shown in Fig. S5 for a reference.

## Discussion

We have investigated color properties of iodine-stained amyloid fibrils of human insulin mainly by using UV-Vis absorbance spectroscopy. When iodine molecules bound on the surface of the insulin fibrils, they showed clear coloration like iodine-starch reactions. An interesting feature of the iodine staining of amyloid fibrils was that different colors were observed among the three types of insulin amyloid fibrils (i.e., NaCl, SDS, and no-salt fibrils) upon the addition of iodine solution. The differences in color tone were spectroscopically demonstrated in visible region, and furthermore, the colors formed by iodine staining could track the propagation of amyloid fibril structures by seed-dependent growth reactions. The present observation suggests that iodine staining can serve as a new probe indicating structural polymorphs of amyloid fibrils.

There are several chemical dyes that have been widely used for detection and tracking of amyloid fibril formation. The most widely used one is ThT, which exhibits enhanced fluorescence upon binding to amyloid fibrils. ThT has been used for many assays as well as extended applications such as microscopic observation of amyloid fibril growth and diagnosis of fibrils in tissue sections^31,52,53^. Several ThT derivatives with high affinity toward amyloid fibrils have also been designed, and recently, radioactively labeled Pittsburgh compound-B is widely used for non-invasive in vivo positron emission tomography imaging for diagnosis of AD^54,55^. Congo Red, which shows characteristic apple-green birefringence under polarized light, is the most histological amyloid dye used for the diagnosis of amyloidosis for several decades^56,57^. 8-Anilino-1-naphthalenesulfonic acid has also been used for detecting fibrils and prefibrillar intermediates, although specificity is not high^58^. These amyloid dyes are very good at detecting the presence and quantifying the amount of amyloid fibrils. However, they are not so sensitive to differences in structure of amyloid fibrils and fail to probe amyloid polymorphism in many cases. Although NaCl, SDS, and no-salt fibrils showed different peak wavelengths in ThT fluorescence spectrum (Fig. 1), ThT fluorescence is typically considered to show little change in peak wavelength and only fluorescence intensity can be used for detecting amyloid polymorphs^59^. Compared to these conventionally used dyes, it is expected that the color formed by iodine staining is highly sensitive to fibril structures, and in the case of the insulin fibril samples used in this study, clear differences in color, which were obvious even with the naked eye, were observed. The present results have indicated that iodine staining can be used for identifying polymorphs of amyloid fibrils.

The coloration of amyloid fibrils by iodine staining is thought to be derived from polymerization of iodine upon binding to amyloid fibrils^10^. In water solution without any host compounds, polyiodide ions larger than I_3_^−^ are absent in an iodine solution^60^. However, a wide range of polyiodides structures have been observed within a supramolecular crystalline compounds and polymers^23–27^. As for amyloid fibrils, it is predicted that, because the cross-β structure of amyloid fibrils has high periodicity along the fibril axis, they can act as a scaffold of iodine complexation in a similar manner to other compounds. Indeed, resonance Raman spectroscopy identified the presence of I_3_^−^ and I_5_^−^ species as fibril-bound forms of iodine (Fig. 5). Considering that I_3_^−^ itself does not absorb visible light so strongly^38,39^, it is predicted that I_5_^−^ and other polyiodide ions composed of I_3_^−^ and I_5_^−^ as fundamental building blocks, (I_3_^−^)_*n*_ and (I_5_^−^)_*n*_, and possibly some other types of polyiodides composed of both of I_3_^−^ and I_5_^−^, (I_3_^−^)_*m*_(I_5_^−^)_*n*_, serve as the molecular basis of color formation.

Interestingly, the resonance Raman spectra excited at 785 nm were similar to the previously reported iodine-amylose pattern in terms of positions and relative intensities of fundamental bands as well as their overtone and combination bands^22,43^. According to the crystal structure of iodine-doped α-cyclodextrin, a structural model that resembles the iodine-amylose complex, it is proposed that a chain-like I_5_^−^ ion is encapsulated inside a channel-like cavity of the helical structure of amylose^27^. It is therefore suggested that linearly aligned (I_5_^−^)_*n*_ polyiodides contribute to the formation of the absorption band at the lowest wavenumber. On the other hand, the band at ~110 cm^−1^ was rather stronger than that at ~160 cm^−1^ when excited at 514.5 nm (Fig. 5a,b), implying that other types of polyiodides containing I_3_^−^, possibly (I_3_^−^)_*n*_, might also be involved in the formation of reddish color. Alternatively, the increased Raman band at ~110 cm^−1^ in the spectra may result from impairment of linear array of (I_5_^−^)_*n*_ chains, in light of a previous report on the analysis of (I_5_^−^)_*n*_ chains with different levels of alignment^44^.

Figure 8 is a proposed model of the binding of polyiodides on amyloid fibrils. The crystal structures of several iodine-doped compounds, such as α-cyclodextrin^21^ and an inorganic-organic compound {(C_4_H_12_N_2_)_2_[Cu^I^I_4_](I_2_)}_*n*_^61^, have already been revealed at atomic level. According to these structures, iodine molecules are encapsulated or buried in gap structures of the host molecules. In contrast, Dzwolak revealed that iodine species bound to bovine insulin fibrils are easily reduced by ascorbic acids^10^, suggesting that iodine molecules are located on the solvent-accessible surface of amyloid fibrils. Given that anisotropic coordination of polyiodides was suggested by the observation of iodine-stained spherulites by polarization microscopy, it is estimated that line shaped polyiodides are formed and stabilized on the grooves of the pleated cross-β structure running in parallel to the long axis of fibrils (model I in Fig. 8). The space between two side chains of the β-strand is approximately 7 Å, which is suitable size for iodine atoms of 4 Å in diameter to get stuck, and it is thus conceived to act as a scaffold to form line shaped polyiodide species. In addition to the surface of cross-β structure, the typical distance between two laminated β-sheet layers is approximately 10 Å. The edge of β-sheets could be another candidate for the binding site of polyiodides in a similar manner to the case of ThT binding^62^ (model II in Fig. 8). Twist grooves formed by intertwining protofilaments may also play a role as iodine binding sites suitable for the fixation of polyiodides. Further investigation is needed for more detailed specification of the binding sites.

**Figure 8.**
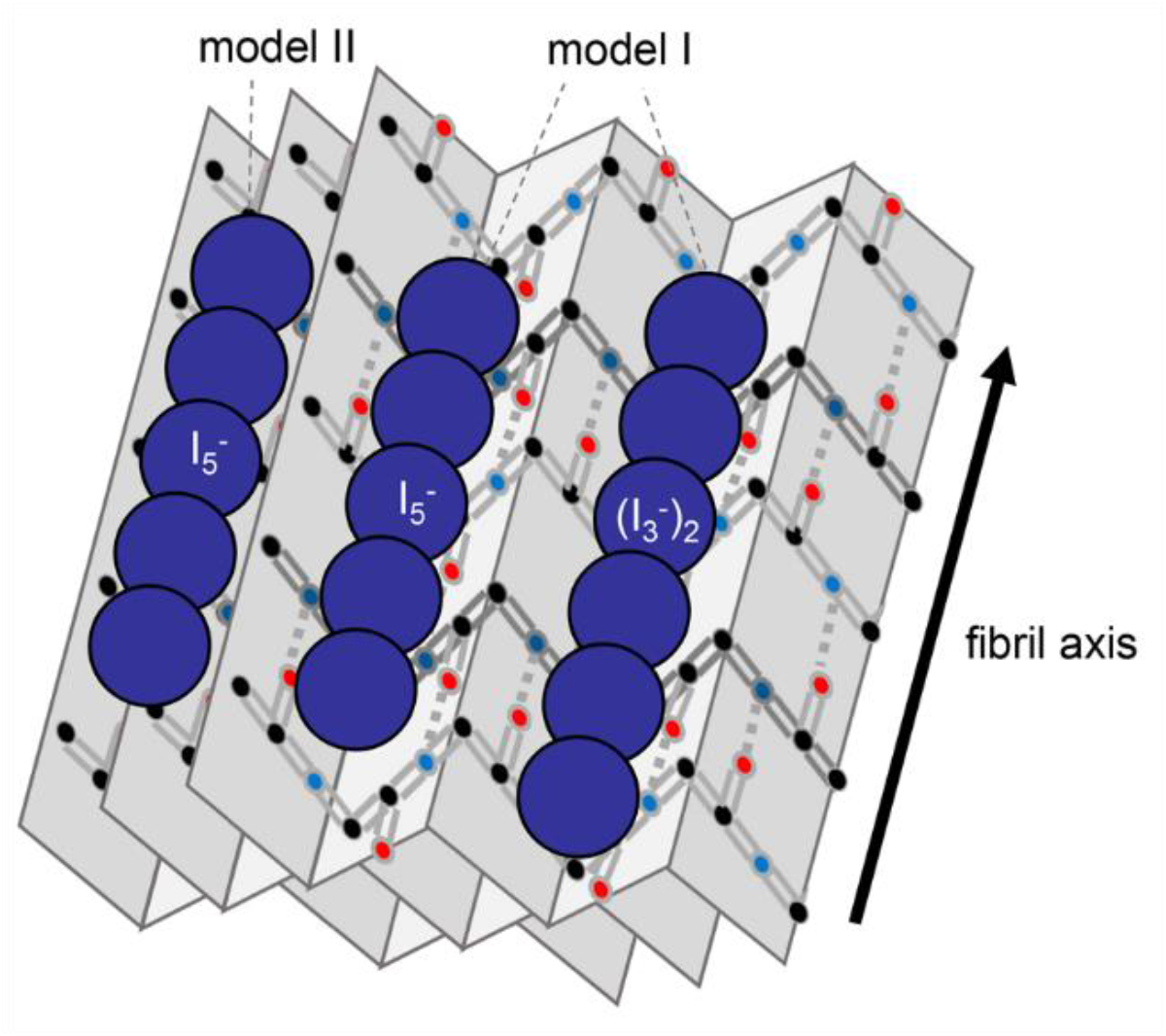
Schematic illustration for the binding of the polyiodides on the surface of amyloid fibrils. In this figure, I_5_^−^ and (I_3_^−^)_2_ are shown as tentative examples of polyiodides, and two possible candidates for the binding sites of polyiodides, i.e., the grooves of the pleated cross-β structure (model I) and the edge of β-sheets (model II), are represented.

In the system of insulin amyloid fibrils, spectral deconvolution of the UV-Vis spectra has clarified that the color formed by iodine staining of amyloid fibrils is represented as the sum of three major absorption bands in visible region. The titration of iodine solution to amyloid fibrils revealed that each of these absorption bands had its own specific binding constant for iodine binding (Fig. 6). This result indicates that insulin amyloid fibrils basically have three independent iodine-binding sites, to each of which a specific and different type of polyiodide ion binds; while the polyiodide ions are commonly composed of I_3_^−^ and I_5_^−^ building blocks, the conformations of the building blocks as well as their composition are different, as deduced from the result of resonance Raman spectra obtained at two different excitation wavelengths (Fig. 5). Additionally, degree of polymerization may also be different depending on the binding sites, in view of previous studies on amylose with different degree of polymerization in which absorption band shifted effectively to longer wavelength as the chain length of polyiodides increased^26^.

As shown in Figure 3, differences in the peak positions and intensity ratio of the three absorption bands in visible region were the spectroscopic entities underlying the color difference in iodine staining among polymorphs. The characteristics of the three absorption bands seem to change sensitively depending on structural properties of fibril surface. It is anticipated that characteristic protein topology of amyloid polymorphs on cross-β structure and/or at its edge parts determines the structure and degree of polymerization of polyiodides on the binding sites, and eventually, leads to different coloration. Furthermore, when the absorption spectra of SDS and no-salt fibrils were compared, the wavenumbers of the three constituent absorption bands were similar to those of SDS fibrils; however, the relative ratio of their absorbance intensities were markedly different (Fig. 3b,c). This observation may suggest that differences in factors related to absorption intensity, i.e., the number of binding site and their binding constant of polyiodides, are also available for detecting polymorphic structures of amyloid fibrils, even though iodine binding sites are very similar.

## Conclusion

Through this work, it was clarified that coloration by iodine staining has promising characteristics in identifying polymorphism of amyloid fibrils. Considering that the colors successfully reported the differences in structure of insulin amyloid fibrils despite that their secondary structures were very similar (Fig. 1–3), it is expected that iodine staining works as a sensitive probe for amyloid polymorphs by discriminating their surface structures. The iodine color also successfully tracked the propagation process of amyloid structures (Fig. 4). Although iodine staining alone cannot function as a technical strategy for clarifying amyloid structures, its colors can act as a fingerprint of amyloid polymorphs. If color information on a particular type of amyloid fibrils of interest is investigated beforehand, its quick identification will be achieved by matching the color features. Moreover, given the recent situation that a growing number of the structures of amyloid fibrils are determined by cryo-EM and ssNMR spectroscopy, valuable insights into the relationships between the surface topology of amyloid fibrils and the corresponding color property by iodine staining are obtained practically by analyzing the color properties of amyloid fibrils with known structures. This will provide perspective on clarifying the molecular mechanism of iodine staining, and consequently, will boost the elucidation of molecular details for amyloid polymorphism underlying amyloid diseases.

Recently, luminescent conjugated polythiophenes and oligothiophenes have been developed as a molecular probe of amyloid fibrils^63^ and very recently, succeeded in differentiate pathological amyloid subtypes of Aβ in AD brains^64^. This observation has revealed the presence of structurally distinct conformations in the affected area of amyloid diseases, arguing the importance of characterizing amyloid polymorphs in biological samples. Although iodine staining is based on absorption and is inferior in terms of signal intensity compared to fluorescent probes, it has potency of being applied to all of the amyloidogenic proteins, and covering a wide range of samples that have not been analyzed so far because of problems with amount that can be supplied for the analysis as well as inhomogeneity. Furthermore, in light of the fact that the iodine staining of amyloid fibrils was firstly discovered in the field of anatomy and is still used in part for diagnosis of amyloidoses, the present work also suggests that it will be worth revisiting the usefulness of the iodine staining of pathological tissues for evaluating the conditions and progress of diseases on the basis of amyloid polymorphism. Further data accumulation connecting the iodine colors and fibril structures will open up new perspectives of the usage of iodine staining in various field of amyloid researches.

## Materials and Methods

### Materials

Recombinant human insulin was purchased from Wako Pure Chemical Industries, Ltd., Osaka, Japan. Insulin was dissolved in 25 mM HCl, and whose concentration was determined using an absorption coefficient of 1.08 for 1.0 mg/ml at 276 nm^65^. Iodine, potassium iodide, and ThT were also purchased from Wako Pure Chemical Industries, Ltd. HCl solution at a concentration of 100 mM, NaCl, and sodium dodecyl sulfate (SDS) were purchased from Nacalai Tesque (Kyoto, Japan).

### Formation of amyloid fibrils of insulin

Human insulin dissolved in 25 mM HCl at a concentration of 2.0 mg/ml was used as a fundamental sample solution. Fibrillation reaction was performed by incubating this solution at 65 °C for 24 hours. For inducing polymorphism of amyloid fibrils, two types of additives, 100 mM NaCl or 100 μM SDS were used in addition to the solvent condition without additives. After the formation of amyloid fibrils, which were referred to as parent fibrils, a seed-dependent fibril formation was performed to obtain daughter fibrils. In this reaction, the fragments of fibrils were added to 2.0 mg/ml insulin dissolved in 25 mM HCl as seeds at a final concentration of 5 % (w/v), and then incubated at 37 °C for 24 hours. It should be noted that all of the seed-dependent fibrillation reactions were performed in the same solvent conditions without any additives. For the preparation of seeds, the parent fibrils were subjected to pulsed sonication using XL2000 sonicator (Misonix, San Diego, CA, USA) operating for 1 sec with power level at 2.0 W. The total number of sonication pulses was 40, and it was ensured that amyloid samples are not excessively heated during the sonication treatment. After the formation of the daughter fibrils, the same seed-dependent fibril formation was repeated using the daughter fibrils as seeds to obtain granddaughter fibrils.

### ThT fluorescence assay

ThT fluorescence assay was performed according to the method of Naiki et al.^53^ with a slight modification of solution; 4.5 μl aliquots of sample solution was mixed with 1.5 ml of 5 μM ThT in 50 mM Gly-NaOH buffer (pH 8.5) at room temperature. After 1 min for equilibrium, fluorescence intensity at 485 nm was measured with an excitation wavelength of 445 nm with RF-5300 PC (Shimadzu Corporation, Kyoto, Japan) to assay the formation of amyloid fibrils.

### Atomic force microscopy (AFM)

Samples of insulin amyloid fibrils were diluted 250 times with 25 mM HCl, and 10 μl aliquots of which were deposited onto a freshly cleaved dry mica surface. The solution was adsorbed for 1 min and then the mica surface was washed with 100 μl of water. The solution was blotted off by blotting paper and the mica plate was air-dried. AFM images were obtained by using SPA-400 and nano navi (Hitachi High-Tech Science, Tokyo, Japan). The scanning tip was a OMCL-AC160TS-C3 micro cantilever (Olympus Corporation, Tokyo, Japan; spring constant = 21-37 N/m, resonance frequency = 270-340 kHz), and the scan rate was 1.0 Hz.

### Attenuated total reflection Fourier transform infrared (ATR-FTIR) absorption spectroscopy

ATR-FTIR spectra were measured with a FT/IR-6100 (JASCO, Tokyo, Japan). 2 μl of sample solution was blotted onto a germanium prism and then dried using compressed air. The spectral resolution and the number of scan were set to 4 cm^−1^ and 128, respectively. A spectrum of atmospheric air was used as a reference spectrum to subtract the contributions of water vapor and carbon dioxide from each sample spectrum.

### Iodine staining of amyloid fibrils

The color formation of insulin fibrils by the addition of iodine was analyzed by using UV-Vis spectroscopy. For the preparation of a sample, iodine solution containing 24 mM KI and 3.2 mM I_2_ in 25 mM HCl was initially prepared, and 0.25 mg/ml insulin fibrils containing 0.3 mM KI, and 0.04 mM I_2_ in 25 mM HCl was made by mixing sample fibrils with the iodine solution. UV-Vis spectra from 250 to 750 nm were then measured at room temperature with a UV-2400PC (Shimadzu Corporation) or Jasco V-650 spectrometer (JASCO Corporation, Tokyo, Japan) with a quartz cell with 1 cm optical length. The reaction between fibrils and iodine molecules was sufficiently rapid to reach equilibrium within 1 min under the present solution conditions, and thus we performed the measurement between 1 and 10 min after preparing the sample. For tracking seed-dependent fibrillation reactions, an aliquot of a reaction mixture was taken every one hour for sample preparation and subsequent spectral measurement, which were performed under the same conditions as those mentioned above. For titration experiments, 1 ml of 0.5 mg/ml insulin fibrils dissolved in 25 mM HCl was prepared as an analyte, to which an iodine solution containing 18 mM KI and 2.4 mM I_2_ in 25 mM HCl was titrated. A UV-Vis spectrum was monitored every after the addition of 4 μl of the iodine solution to the fibrils, and this procedure was repeated until spectral saturation was observed.

### Resonance Raman spectroscopy

Raman resonance spectra were measured by using two types spectrometers, HR-800-LWR (HORIBA, Kyoto, Japan) with 514.5 nm excitation and inVia Raman microscope (Renishaw, Gloucestershire, UK) with 785 nm excitation. Both of the spectrometers were equipped with notch filters for suppressing Rayleigh scattering. A sample containing 0.5 mg/ml insulin fibrils, 0.6 mM KI, and 0.08 mM I_2_ in 25 mM HCl was taken in a glass capillary with its inner diameter of 1.13 mm (Drummond Scientific Company, Broomall, PA, USA) and placed on a sample stage. An excitation light was focused on the sample solution closed to the wall surface of the capillary to minimize reabsorption of Raman scattered light by the sample. Raman scattering was measured with back-scattering mode from 0 to 500 cm^−1^ at room temperature.

### Polarization microscopy of spherulites

To investigate polarization properties of the polyiodide ions bound on amyloid fibrils, microscopy observation of iodine-stained insulin spherulites was performed with an ECLIPSE LV100POL (Nikon, Tokyo, Japan), in which an analyzer was placed in the optical pathway between an objective lens and an eyepiece. To prepare spherulites, insulin solution at a concentration of 5.0 mg/ml in 25 mM HCl was sealed between slide and cover glasses and incubated at 70 °C for 2 hours, according to the method reported by Rogers et al^51^. The spherulites formed were then stained with iodine by adding a small amount of iodine solution containing 24 mM KI and 3.2 mM I_2_ in in 25 mM HCl. Microscope images were recorded with a DS-L3 digital camera (Nikon) at different angles of rotations of the analyzer.

## Data availability

All data generated or analysed during this study are included in this published article (and its Supplementary Information files).

## Acknowledgments

We thank Tomomi Kozu (Renishaw K.K.) for help of resonance Raman measurements and Prof. Masahide Yamamoto (Kyoto University) for valuable comments on iodine-polymer complexes. AFM measurements were performed at Research Facility Center for Science and Technology, Kobe University. This work was supported by JSPS Core-to-Core Program, A. Advanced Research Networks. This work was funded by JSPS KAKENHI Grant Numbers JP16H04778, JP16H00772, and JP17H06352.

## Additional information

Supplementary information, which contains 5 supplementary figures, is attached to this manuscript.

## Notes

### Competing Interest Statement

The authors have declared no competing interest.

