## Supporting Information for "Iodine staining as a useful probe for distinguishing amyloid fibril polymorphs"

**Figure S1.** UV-vis spectra showing negligible interaction between iodine molecules and native insulin.

**Figure S2.** UV-vis spectra showing negligible effects of the presence of salts on iodine staining.

**Figure S3.** Representative results of curve fitting of the UV-vis absorption spectra in the seeding reaction.

**Figure S4.** Representative results of curve fitting of the UV-vis absorption spectra in the iodine titration experiment.

**Figure S5.** Images of an unstained spherulite observed by polarization microscopy.

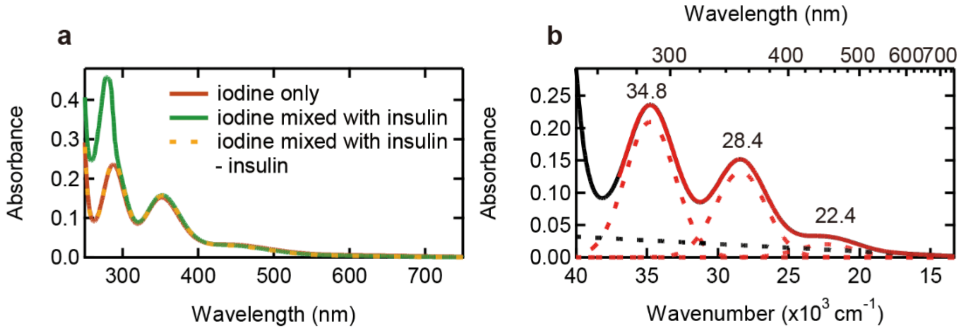


**Figure S1. UV-vis spectra showing negligible interaction between iodine molecules and native insulin.** (**a**) UV-vis spectra of iodine solution (red line), iodine solution mixed with native insulin (green line), and a differential spectrum obtained by subtracting the spectrum of native insulin from that of the iodine solution mixed with native insulin (dashed orange line). The iodine staining was performed under the conditions of 0.3 mM KI and 0.04 mM $I_2$ in 25 mM HCl, and the concentration of native insulin was 0.25 mg/ml. The dashed orange line was overlapped with the red line, suggesting that there was no significant interaction between iodine and native insulin. (**b**) Result of curve fitting of the spectrum of iodine solution. A black line represents the experimental spectrum, and a red line represents the fitted spectrum using eq. 1 assuming three Gaussian bands (i.e., $n$ = 3). The Gaussian bands and the baselines are represented with dashed red lines and a dashed black line, respectively.

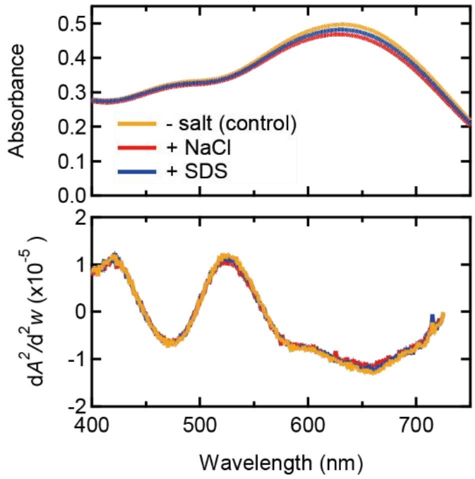


**Figure S2. UV-vis spectra showing negligible effects of the presence of salts on iodine staining.** Absorption spectra in visible region (top) and their second derivative spectra (bottom) of iodine-stained no-salt fibrils (orange) and those monitored in the presence of NaCl (red) or SDS (blue). No-salt fibrils were initially prepared, and after the treatment with ultrasonic pulses, NaCl or SDS was added to the fibrils at a final concentration of 5 mM or 5 μM, respectively; these concentrations correspond to those of NaCl or SDS contained in the NaCl/SDS daughter fibrils coming from the parent seed fibrils. After the incubation of the amyloid fibrils at 37 °C for 24 hours, iodine staining was performed at final concentrations of 0.25 mg/ml amyloid fibrils, 0.3 mM KI, and 0.04 mM $I_2$ in 25 mM HCl. The observed spectra were almost unchanged even after the addition of NaCl or SDS, eliminating the effect of the residual salts on the color formation by iodine staining.

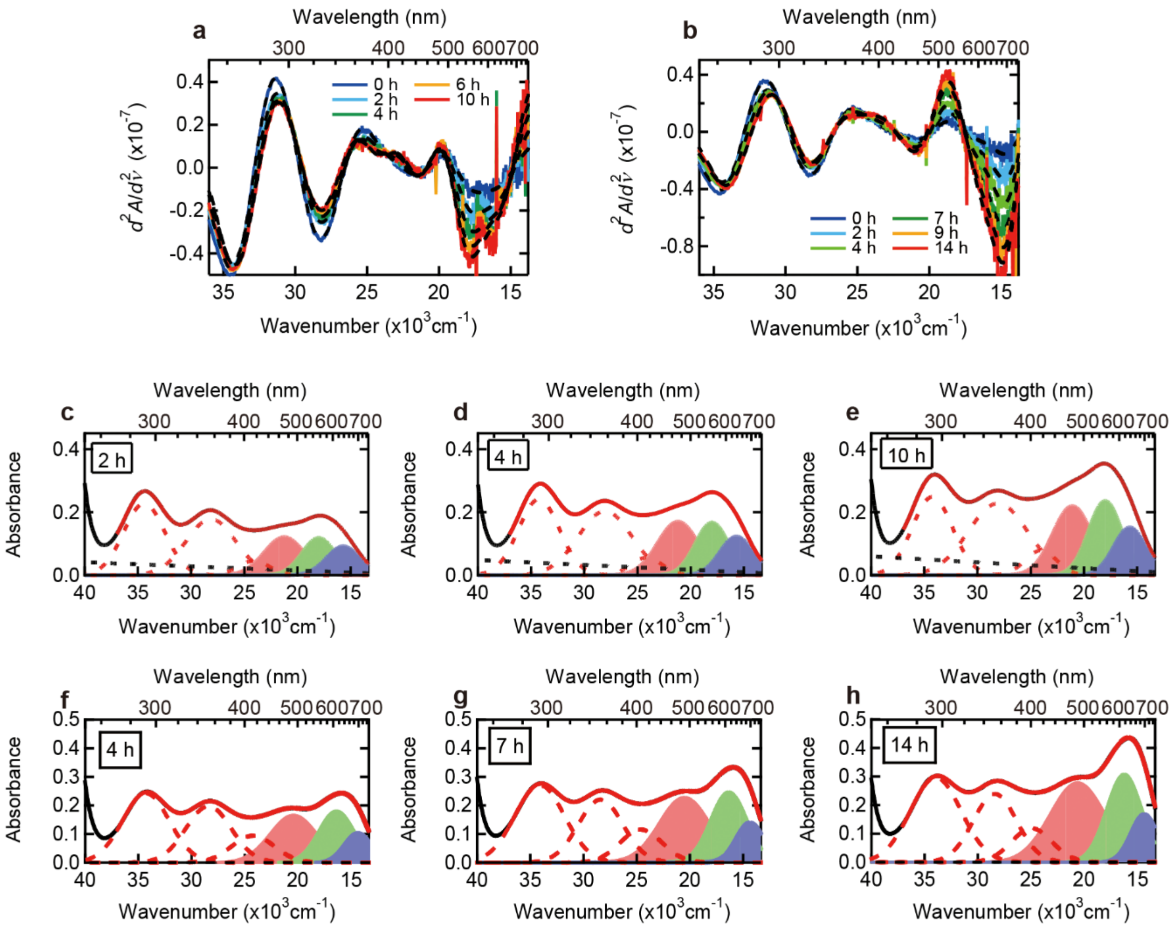


**Figure S3. Representative results of curve fitting of the UV-vis absorption spectra in the seeding reaction.** (**a** and **b**) Second derivatives of the absorption spectra of iodine-stained NaCl (**a**) and SDS fibrils (**b**) shown in Fig. 4a and c. Dashed lines are the second derivatives of the fitted spectra obtained by the curve fitting with six Gaussian peaks, which was constructed to assess the validity of the curve fitting. (**c**-**h**) Results of curve fitting of NaCl (**c**-**e**) and SDS fibrils (**f**-**h**) at three representative time points. In each panel, black and red solid lines represent the experimental and the fitted spectra, respectively. Six Gaussian peaks are shown with red dashed lines or filled areas and a baseline is shown with a black dashed line. In performing curve fitting, the position and FWHM of the band II (green filled area) for NaCl fibrils or the band III (blue filled area) for SDS fibrils, the most prominent band among the three bands with the largest minimum in the second derivatives, were fixed to values read from the results of second derivatives.

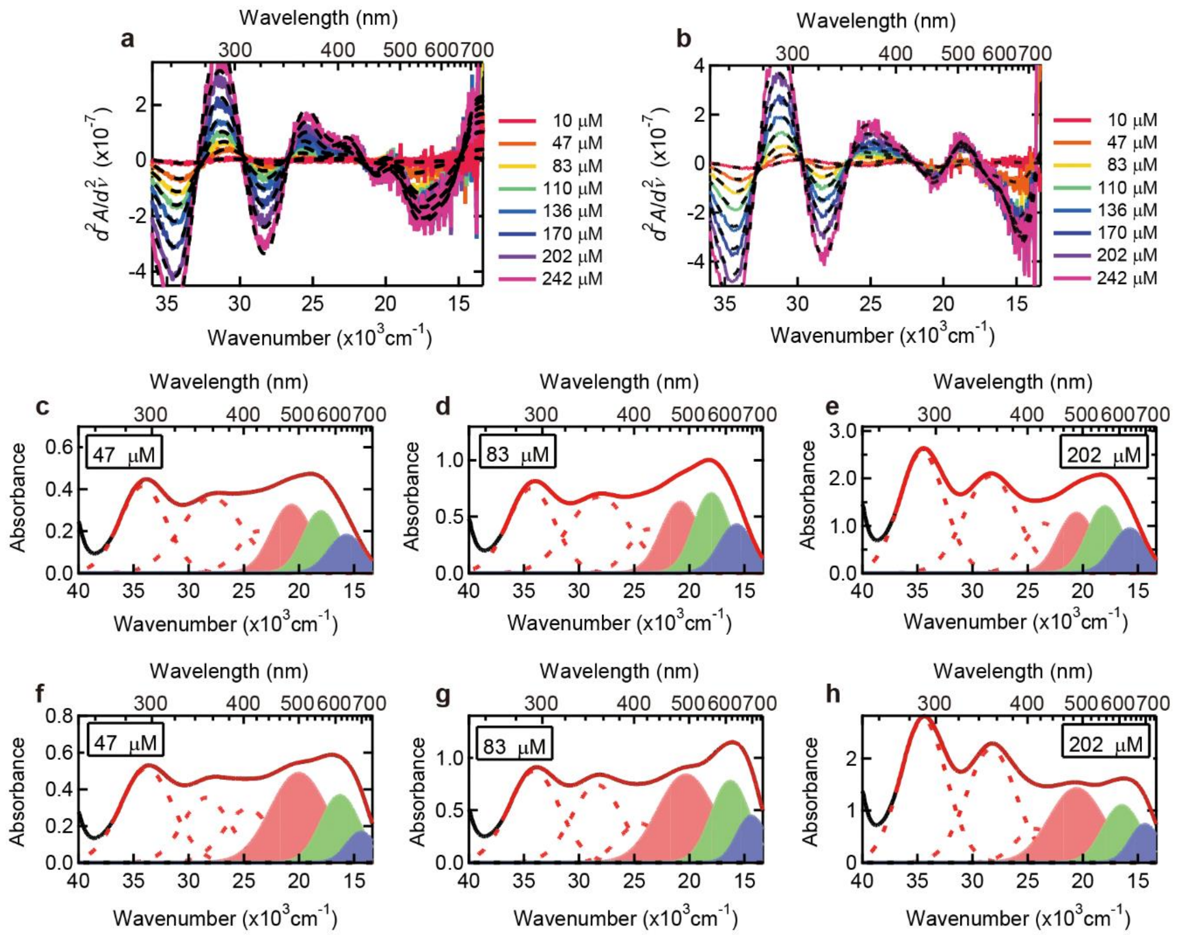

**Figure S4. Representative results of curve fitting of the UV-vis absorption spectra in the iodine titration experiment.** (**a** and **b**) Second derivatives of the absorption spectra of iodine-stained NaCl (**a**) and SDS fibrils (**b**) shown in Fig. 6a and b. Dashed lines are the second derivatives of the fitted spectra obtained by the curve fitting with six Gaussian peaks, confirming the consistency of the fitting results. (**c**-**h**) Results of curve fitting of NaCl (**c**-**e**) and SDS fibrils (**f**-**h**) at three representative iodine concentrations. In each panel, black and red solid lines represent the experimental and the fitted spectra, respectively. Six Gaussian peaks are shown with red dashed lines or filled areas and a baseline is shown with a black dashed line. In performing curve fitting, the position of the band II (green filled area) for NaCl-fibrils or the band III (blue filled area) for SDS-fibrils and its FWHM were fixed based on the results of second derivatives.

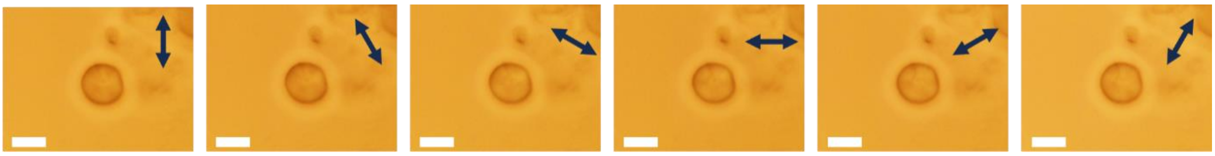

**Figure S5. Images of an unstained spherulite observed by polarization microscopy.** The direction of the analyzer is represented with a double-headed arrow in the upper right of each image. Scale bars represent 10 μm.
